# The effects of iron deficient and high iron diets on SARS-CoV-2 lung infection and disease

**DOI:** 10.1101/2024.05.29.596393

**Authors:** Agnes Carolin, David Frazer, Kexin Yan, Cameron R. Bishop, Bing Tang, Wilson Nguyen, Sheridan L. Helman, Jay Horvat, Thibaut Larcher, Daniel J. Rawle, Andreas Suhrbier

**Affiliations:** Inflammation Biology, QIMR Berghofer Medical Research Institute, Brisbane, Queensland, 4029, Australia; Molecular Nutrition, QIMR Berghofer Medical Research Institute, Brisbane, Queensland, 4029, Australia; School of Biomedical Sciences and Pharmacy, Faculty of Health and Medicine, University of Newcastle and Hunter Medical Research Institute, Australia; UMR0703 APEX, INRAE, Oniris, Nantes CEDEX 03, France; GVN Centre of Excellence, Australian Infectious Disease Research Centre, Brisbane, Queensland 4029 and 4072, Australia

**Keywords:** : Iron deficiency, iron loading, SARS-CoV-2, omicron XBB, C57BL/6J mice, inflammation, lung, RNA-Seq

## Abstract

The severity of Coronavirus disease 2019 (COVID-19) caused by the severe acute respiratory syndrome coronavirus 2 (SARS-CoV-2) is often dictated by a range of comorbidities. A considerable literature suggests iron deficiency and iron overload may contribute to increased infection, inflammation and disease severity, although direct causal relationships have been difficult to establish. Here we generate iron deficient and iron loaded C57BL/6J mice by feeding low and high iron diets, with mice on a normal iron diet representing controls. All mice were infected with a primary omicron XXB SARS-CoV-2 isolate and lung inflammatory responses were analyzed by histology, immunohistochemistry and RNA-Seq. Compared with controls, iron deficient mice showed no significant changes in lung viral loads or histopathology, whereas, iron loaded mice showed slightly, but significantly, reduced lung viral loads and histopathology. Transcriptional changes were modest, but illustrated widespread dysregulation of inflammation signatures for both iron deficient vs. controls, and iron loaded vs. controls. Some of these changes could be associated with detrimental outcomes, whereas others would be viewed as beneficial. Diet-associated iron deficiency or overload thus induced modest modulations of inflammatory signatures, but no significant histopathologically detectable disease exacerbations.

**Author summary:** A diet deficient in iron can lead to anemia, a widespread problem worldwide. A diet with excessive iron is less common, but can be associated with excessive consumption of iron supplements. We investigate herein using a mouse model, whether low or high iron diets predispose to detrimental outcomes in the lungs after infection with SARS-CoV-2. A considerable literature suggests iron dysregulation would promote infection and inflammation. However, we found, although inflammatory responses showed modest modulations, viral loads were unaffected or slightly reduced, and lung histopathology was either unaffected or indicated slightly less severe disease. These findings do not support a view that low or high iron diets represent comorbidities predisposing to overt detrimental outcomes for acute COVID-19 lung disease.

## Introduction

The severe acute respiratory syndrome coronavirus 2 (SARS-CoV-2) is the etiological agent of Coronavirus disease 2019 (COVID-19) [1, 2] and has caused a global pandemic involving ∼775 million cases and ∼7 million deaths worldwide [3]. COVID-19 is often associated with a ‘cytokine storm’ and the life threatening, acute respiratory distress syndrome (ARDS). The severity of COVID-19 is influenced by a range of comorbidities [4–6], with a large body of literature suggesting that iron deficiency and iron overload may also contribute to disease severity [7–19].

In mammals hundreds of proteins use iron in a multitude of cellular activities [20] including inflammation and immunity [21]. Iron levels and distributions in different tissues and cells, under different conditions, are regulated by a complex network of processes [22]. For instance, multiple studies have shown that SARS-CoV-2 infection disrupts iron homeostasis and/or modulates iron-associated biomarkers [10, 14, 23–32]. This is not unique to SARS-CoV-2 as many infections modulate iron biomarkers [33], with clinical determinations of iron status thus generally unreliable in patients presenting with infectious and/or inflammatory diseases [34, 35]. Establishing the iron status of a patient presenting with COVID-19 is thus difficult. How patients’ iron status prior to SARS-CoV-2 infection affects the severity of COVID-19 after infection is often difficult to explore in clinical settings, as patients tend to present only after they have developed disease.

Iron deficiency is a widespread problem and is associated with a range of clinical issues, primarily anemia [36, 37]. Pre-existing anemia has been associated with increased mortality risk for hospitalized COVID-19 patients [11], perhaps due to SARS-CoV-2 infection further exacerbating the anemia [38]. Dietary iron deficiency is responsible for about half the 2 billion cases of anemia globally [36, 39]; however, anemia can have a variety of causes; globally this primarily involves thalassemias, sickle cell trait, and malaria [40]. Other conditions are also associated with anemia, including alcoholism [41], diabetes [42], cardiovascular disease [43] and chronic obstructive pulmonary disease [44]. Whether anemia arising from an iron deficient diet, or the comorbidity giving rise to the anemia, is responsible for the increase in COVID-19 severity remains unclear [45–49]. Iron dysregulation and inflammatory stress erythropoiesis have also been associated with long-COVID [12]. However, cause and affect are again difficult to verify, as increased iron dysregulation and stress erythropoiesis may be the result of more severe SARS-CoV-2 infections [38, 50], which then predispose to more pronounced long-COVID [51, 52].

A number of publications have speculated on a connection between iron overload and increased severity of COVID-19, largely based on iron biomarker studies [9, 13–18]. In addition, iron loaded mice, injected with a pseudovirus dislaying the spike protein, showed an increase in serum CCL4, IL1β, IL-6 and TNFα levels [8]. Iron overload can arise from a number of conditions, most famously hereditary hemochromatosis or thalassemias [53], which lead to a unique pattern of body and cellular iron distributions [54, 55]. However, iron overload can also arise from excessive dietary iron intake, which is primarily associated with excessive iron supplement consumption [56–59], but can also be associated with a high iron diet [60]. Whether pre-existing iron loading due to a high iron diet (in the absence of co-morbidities) exacerbates acute COVID-19 severity remains largely unexplored.

Here we use adult wild-type C57BL/6J mice and established models of diet-induced iron deficiency and iron overload [61–64], and SARS-CoV-2 infection [65, 66]. Iron deficient and iron loaded mice were compared with control mice (fed a normal iron diet) after infection with a primary human omicron XBB isolate of SARS-CoV-2 [67]. The impacts of these diets on viral replication and inflammatory disease in the infected lung were characterized using histology (H&E staining), immunohistochemistry and RNA-Seq at 2 days post infection (dpi) (peak viral load) and 6 dpi (peak of acute immune pathology) [65, 66]. Although widespread modulations in transcriptional signatures associated with inflammatory responses were observed, neither iron deficiency nor iron overload resulted in significant increases in viral replication or histopathology.

## Results

### Iron deficient vs. control diet; weights, lung viral loads and liver iron levels

Male C57BL/6J mice were fed either a control diet (normal iron) or a diet deficient in iron for 7 weeks (Fig. 1a). The iron deficient diet resulted in a significant reduction in growth rates so that mice were ≈5 grams lighter just prior to infection (0 dpi) (Fig. 1b). XBB infection had no significant effects on the weight of mice in either the control or the iron deficient groups (Supplementary Fig. 1). Lungs were harvested on 2 dpi (peak viral titers) and 6 dpi (day of peak lung pathology) [66, 68]. There were no significant differences in lung tissue titers (Fig. 1c), or in lung viral read counts as determined by RNA-Seq (Fig. 1d). Viral titers in the nasal turbinates were slightly, but significantly, lower (0.62 log_10_CCID_50_/g, p=0.009) at 2 dpi in mice fed the iron deficient diet (Supplementary Fig. 2a). Liver iron levels were quantitated, with mice on the iron deficient diet showing significantly lower liver iron levels (Fig. 1e), confirming that this diet had successfully reduced iron levels.

**Fig. 1.**
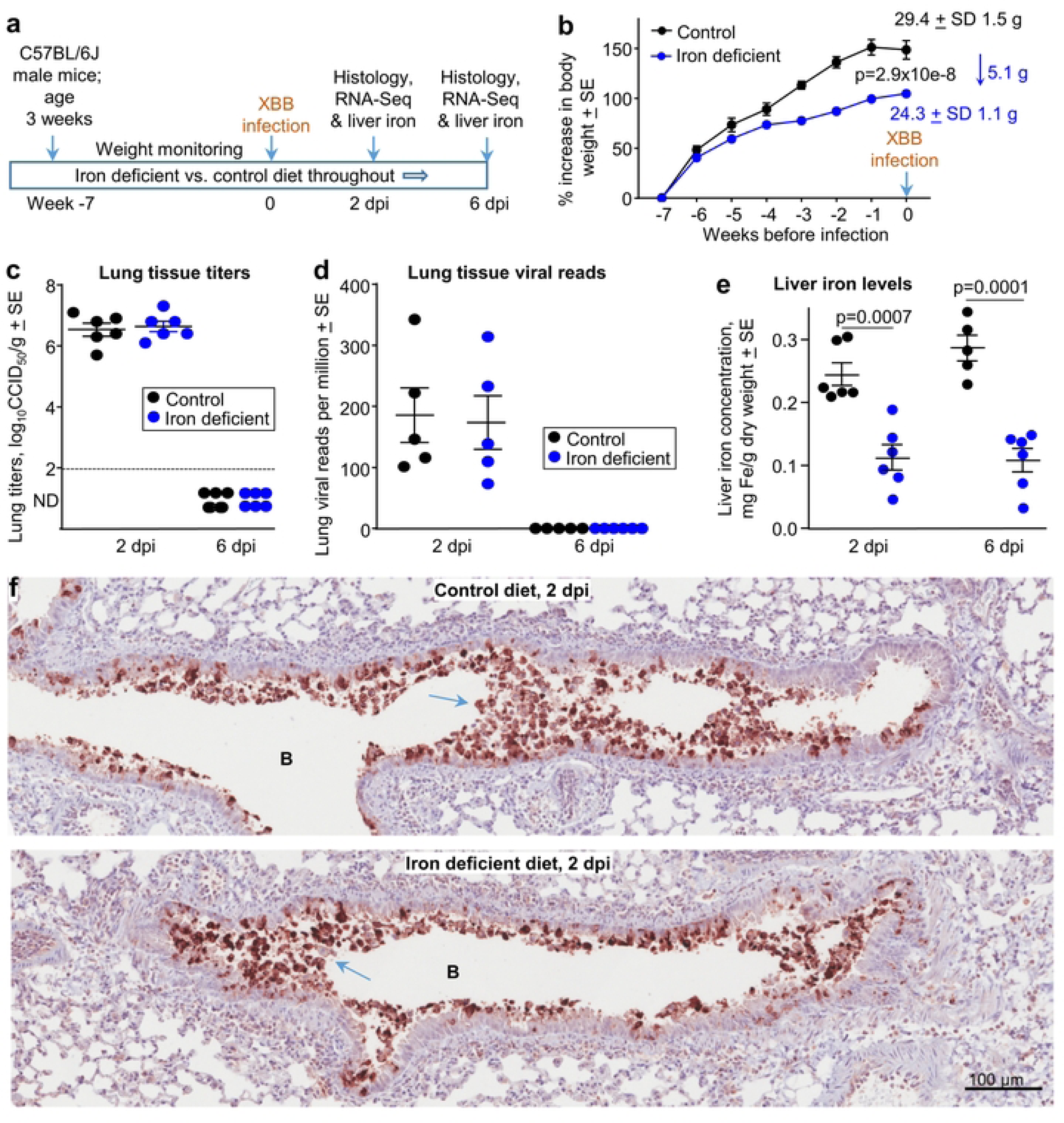
Iron deficient vs. control diets; weights, viral loads and iron levels. **a** Time line of experiment. **b** Mean percentage increases in mouse body weights prior to XBB infection. Mean body weights ± SD in grams are also provided at week 0 (also 0 dpi). Statistics by t test for weight differences at week zero, n=11/12 mice per group. There was no significant weight changed after infection (Supplementary Fig. 1). **c** Lung tissue XBB virus titers. Limit of detection ≈2 log_10_CCID_50_/g (dashed line); ND – not detected. **d** XBB viral read counts obtained from RNA-Seq of lung tissues. **e** Liver iron levels in the same mice shown in b-d. Legend as in d. Statistics by t tests. **f** IHC of mouse lung at 2 dpi for mice on the control diet compared with mice on an iron deficient diet. B – bronchial air space. Blue arrows **-** sloughing of virus infected bronchial epithelial cells (and associated cell debris) into the bronchial lumen.

Thus, although an iron deficient diet significantly reduced the growth of the mice, SARS-CoV-2 tissue titers were not significantly different in lungs, and were slightly lower in nasal turbinates.

### Iron deficient vs. control diet; immunohistochemistry

Whole lung sections were stained with a SARS-CoV-2 specific monoclonal antibody [69]. Staining was primarily associated with the bronchial epithelium and cellular debris in the bronchial lumen (Fig. 1f). The latter likely represents bronchial epithelial cells sloughed-off into the airways after infection-induced cytopathic effects (CPE). No overt differences in staining was observed for mice on the iron deficient vs. control diets, consistent with data in Fig. 1c,d. IHC staining of an uninfected mouse lung, illustrating the low level of background staining, is shown in Supplementary Fig. 3a.

### Iron deficient vs. control diet; histochemistry and histopathology

Whole lung sections were stained with H&E and scanned slides were examined by a European board-certified veterinary pathologist for histopathological lesions. Lung lesions were scored using 6 criteria with examples shown; (i) emphysema (Fig. 2a), (ii) bronchial epithelium damage (Fig. 2b), (iii) bronchial content (Fig. 2c), which occasionally included red blood cells (RBC) (Supplementary Fig. 3b), (iv) vascular changes comprising leukostasis (Fig. 2d,e), perivascular hemorrhage (Fig. 2d) and/or leukocytoclasis (Fig. 2f), (v) perivascular edema (Fig. 2g) and (vi) perivascular and/or peribronchial cuffing (Fig. 2h). Scores were summed to provide a cumulative score for each mouse (Supplementary Fig. 4a), with no significant differences emerging between groups (Fig. 2i).

**Fig. 2.**
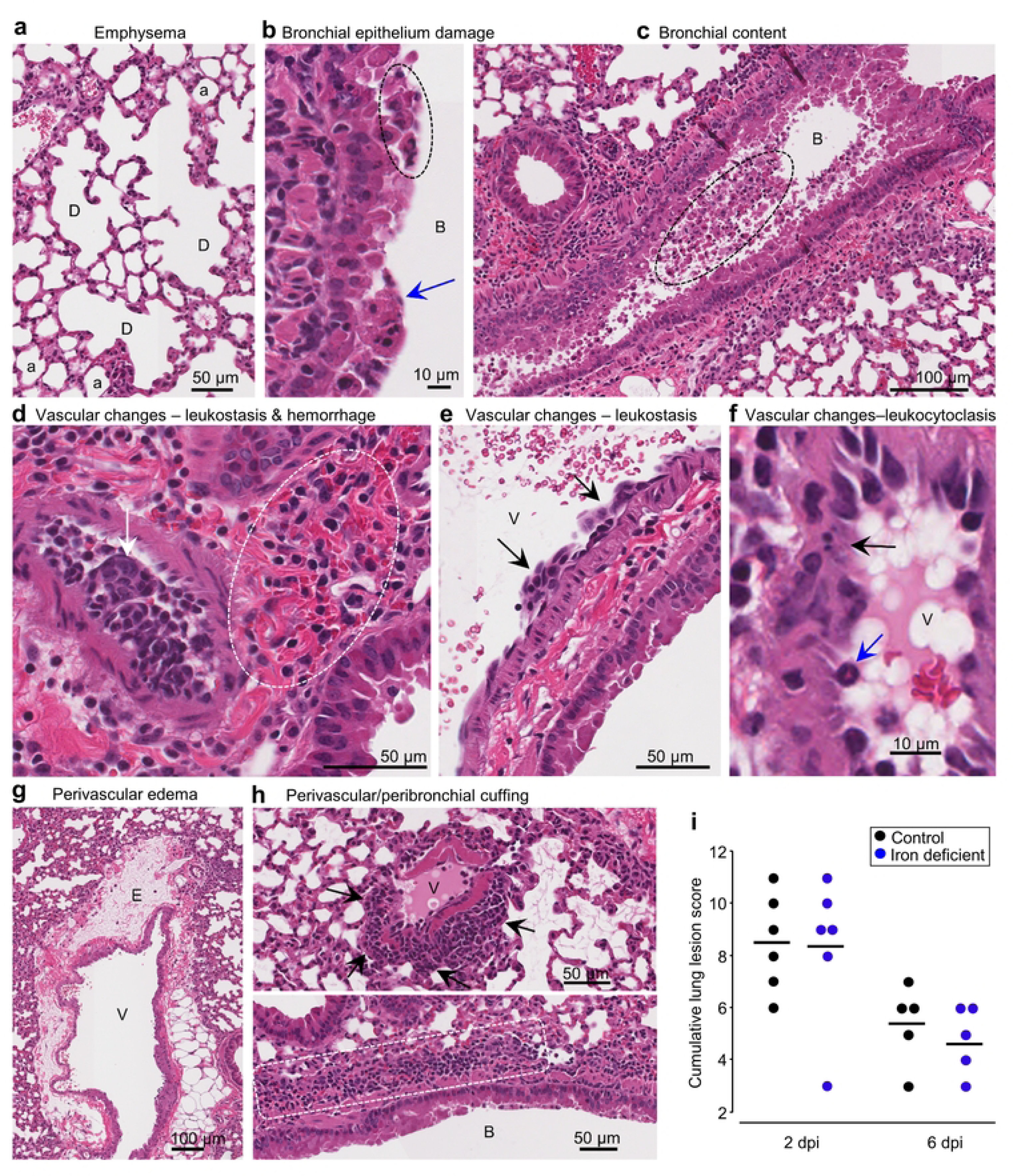
Iron deficient vs. control diets; lung histopathology. H&E staining of infected lungs. **a** Iron deficient diet 2 dpi showing dilated coalescing alveoli (D) illustrating emphysema. a – alveoli. Whole slide score for emphysema = 1 on a 0-2 scale. **b** Control diet 2 dpi showing necrotic bronchial epithelia cells (blue arrow) and partial loss of bronchial epithelial architecture (dashed oval). B – bronchial lumen. Whole slide score for bronchial epithelial damage = 2 on a 0-3 scale. **c** Control diet 2 dpi showing cellular debris in bronchial air space (dashed oval); this material stains positive for viral antigen. B – bronchial air space. Whole slide score for bronchial content = 2 on a 0-3 scale. **d** Control diet 2 dpi showing occlusion of the vascular lumen by clumps of leukocytes (leukostasis) (white arrow), with perivascular hemorrhage (dashed oval). Whole slide score for vascular wall changes = 3 on a 0-3 scale. **e** Iron deficient diet 6 dpi showing clumps of leukocytes adherent to the blood vessel intima (arrows). V – vascular lumen. Whole slide score for vascular wall changes = 2. **f** Control diet 2 dpi showing leukocytoclasis (vascular damage caused by nuclear debris from infiltrating neutrophils) (black arrow). Blue arrow – neutrophil. Whole slide score for vascular wall changes = 3. **g** Control diet 2 dpi showing perivascular edema (E). V – vascular lumen. Whole slide score for edema = 2. **h** (Top) Iron deficient diet 2 dpi showing perivascular cuffing (arrows). V – vascular lumen. Whole slide score for perivascular/peribronchial cuffing =2. (Bottom) Control diet showing peribronchial cuffing (white dashed box showing leukocytes). B – bronchial lumen. Whole slide score for perivascular/peribronchial cuffing =3. **i** Cumulative lung lesion scores for each mouse; raw data is shown in Supplementary Fig. 4a.

White space analysis, an approximate measure of lung consolidation, showed the expected [70–72] significant reduction in virus-infected mice compared with uninfected mice (Supplementary Fig. 5). However, no significant differences emerged between mice on the different diets, although white space reductions appeared to have occurred slightly earlier (2 dpi) in some iron deficient mice (Supplementary Fig. 5).

Using the same H&E stained sections, a pixel count analysis was undertaken to generate a ratio of nuclear (purple) to cytoplasmic (red) staining, which provides an approximate measure of leukocyte infiltration [72, 73]. As expected, infected mice showed significantly more infiltrates than naïve mice; however, no significant differences emerged between infected mice on the different diets (Supplementary Fig. 6a,b).

### Iron deficient vs. control diet; lung RNA-Seq at 2 dpi

RNA-Seq analysis of lungs at 2 dpi from mice fed an iron deficient vs. a control diet identified only 109 DEGs, with all but 5 of these showing low (<1 log_2_) fold change (Supplementary Table 1; PC2/PC1 plots are shown in Supplementary Fig. 7a). A heat map of the top 100 genes that provided the greatest contribution to the segregation between groups, further illustrated that gene expression differences between the groups was low and not particularly focused on any specific set of genes (Supplementary Fig. 8a,b). The results from the bioinformatic analyses are shown in Supplementary Table 1 and are summarized in Fig. 3a (see below).

**Fig. 3.**
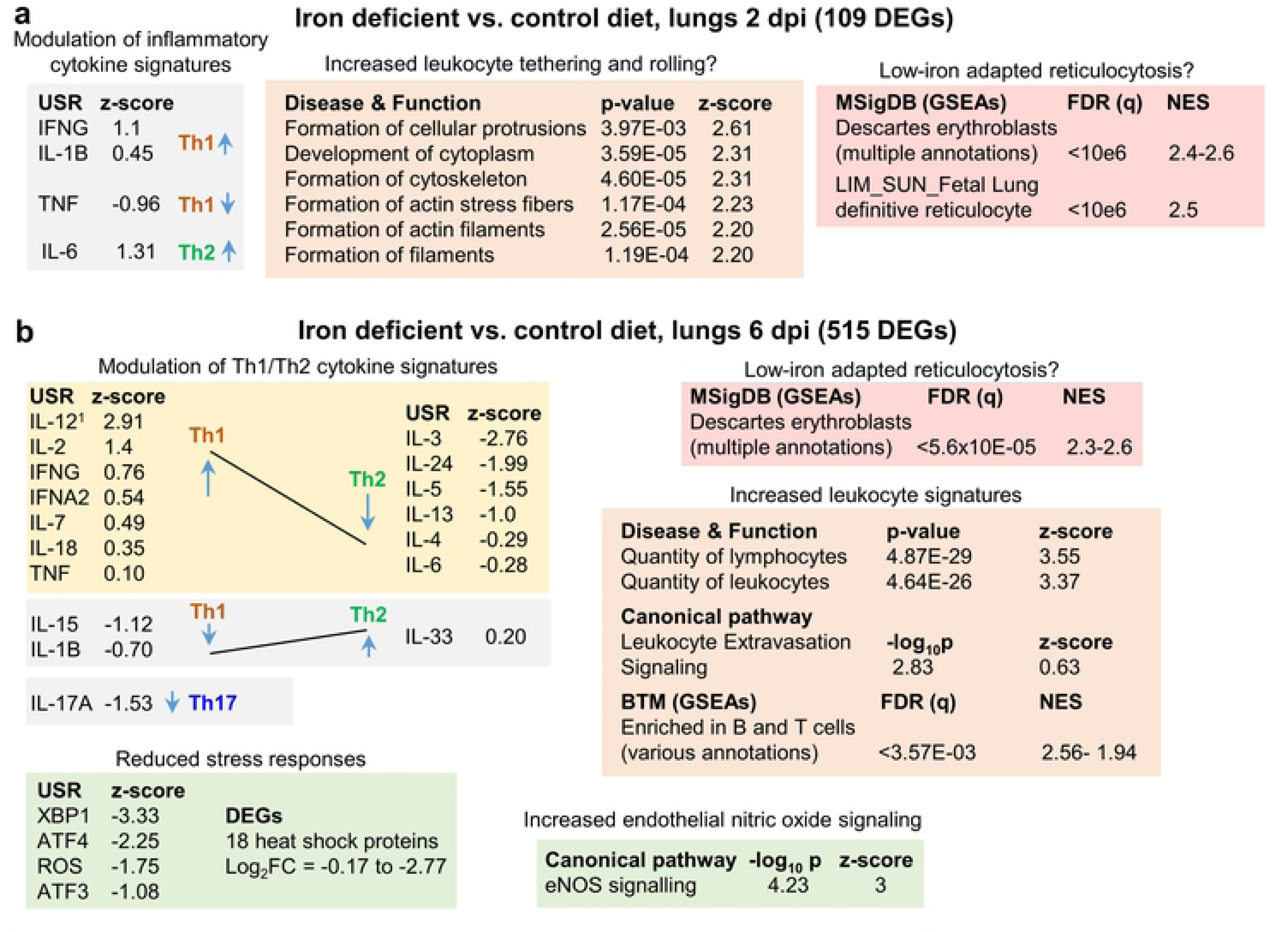
Iron deficient vs. control diets; bioinformatic summary. **a** RNA-Seq analysis of lungs from mice fed an iron deficient vs. control diet at 2 dpi yielded 109 DEGs. DEGs were analysed by IPA using the Up-Steam Regulator (USR) (cytokine annotations shown), and Disease and Function features. The All genes list was interrogated using GSEAs and gene sets provided by MSigDB. Full data sets are provided in Supplementary Table 1. ^1^USR annotation is “IL-12 (family)” and Molecular Type is “group” rather than “cytokine”. **b** RNA-Seq analysis of lungs from mice fed an iron deficient vs. control diet at 6 dpi yielded 515 DEGs. Bioinformatic summary as for a, with the addition of USRs associated with stress responses, and IPA Canonical pathway annotations. Full data sets are provided in Supplementary Table 2.

The 109 DEGs were analyzed by Ingenuity Pathway Analysis (IPA). Using the Up Stream Regulator (USR) feature, mild modulation of inflammatory cytokine signatures was identified (Fig. 3a). As iron deficiency has been associated with changes in the Th1/Th2 cytokine balance [21, 74, 75], the Th1 or Th2 association for each cytokine is shown (Fig. 3a), although 2 dpi is generally too early for significant adaptive T cell responses. The IPA Diseases or Functions feature identified a series of top annotations associated with cellular protrusions, cytoskeleton and actin (Fig. 3a). These annotations may indicate increased leukocyte tethering and rolling [76] in the iron deficient mice. However, significant histological differences were not apparent (Supplementary Fig. 9), as might be expected with only 109 DEGs.

Gene set enrichment analyses (GSEAs) were undertaken using the ‘All genes’ list (ranked by fold change) (Supplementary Table 1) and gene sets from the Molecular Signatures Database (MSigDB) [77, 78]. This analysis identified a series of annotations, with high Normalized Enrichment Scores (NES), that were associated with erythroblasts (Fig. 3a, NES ≈ 2.4-2.6). Erythroblasts are ordinarily restricted to bone marrow, with the loss of the nucleus from these cells preceding release of the resulting reticulocytes, immature red blood cells (RBC), into the circulation. Reticulocytes retain the mRNA profile of erythroblasts in the final stages of maturation [79]. These erythroblast annotations thus likely reflect an increase in gene signatures associated with reticulocytosis in iron deficient mice. Viral infections, including COVID-19, can result in significant damage to RBC [80, 81], with COVID-19 also able to infect RBC progenitors [82]. Erythropoiesis is thus stimulated [12], with erythroblasts adapting to iron deficient conditions by modulating gene expression [83–85], with such modulation likely giving rise to these erythroblast annotations.

In summary, at 2 dpi iron deficiency imparted only mild transcriptional changes, which were associated with minor modulation of inflammatory cytokine, and perhaps tethering and rolling, signatures. Erythroblast/reticulocyte signatures also indicated erythroblast adaptation to low iron conditions.

## Iron deficient vs. control diet; lung RNA-Seq at 6 dpi

RNA-Seq analysis of lungs at 6 dpi from mice fed an iron deficient diet vs. a control diet identified 515 DEGs, with all but 23 of these showing low (<1 log_2_) fold change (Supplementary Table 2). The PC2/PC1 plot is shown in Supplementary Fig. 7a. The results from the bioinformatic analyses are shown in Supplementary Table 2 and are summarized in Fig. 3b (see below).

IPA USR analysis again identified modulation of cytokine response signatures that were generally associated with up-regulation of Th1 signatures and down-regulation of Th2 signatures (Fig. 3b). This contrasts with previous reports suggesting immune activation under iron-deficient conditions results in the expansion of Th2, but not Th1 cells [74, 75], although such Th1/Th2 modulation is likely to be setting dependent [21]. For instance, iron deficiency is reported to blunt IL-6 responses [86, 87] in some settings, but not others [88]. Consistent with the observations herein (Fig. 3b), iron deficiency has been reported to reduce IL-4 [89] and IL-17A responses [90, 91]. In addition, iron deficiency has been reported to cause non-proliferating, altruistic T cells to produce IL-2 [92].

Top IPA Diseases or Functions annotations indicated an increase in the quantity of lymphocytes/leukocytes in infected iron-deficient mouse lungs (Fig. 3b; Increased leukocyte signatures). BTM GSEAs suggest these increases are primarily associated with B cells (Supplementary Table 2), consistent with identification of CXCR5 as an upregulated DEG (log_2_FC=0.88) (Supplementary Table 2). CXCR5 is the receptor for CXCL13, which is the key chemokine for B cell recruitment to sites of inflammation [93, 94]. However, histological analyses indicated only marginal, non-significant, increases in leukocytes in iron deficient mice (Supplementary Fig. 6a, 6 dpi; Supplementary Fig. 4a, cuffing 6 dpi), arguing the transcriptional modulations (Fig. 3b) do not reflect an overall overt increase in inflammatory infiltrates. Instead, they may reflect transcription changes associated with (i) modest increases in infiltrates, with, for instance, the extravasation annotation indicating a relatively low z-score (Fig. 3b, z-score = 0.63), and/or (ii) Th1/Th2 modulation changing leukocyte transcriptional profiles, and/or (iii) changes in the type of cells infiltrating the infected lungs in iron deficient mice (see below).

The top upregulated Canonical pathway was endothelial nitric oxide synthase (Fig. 3b, eNOS), with increased endothelial nitric oxide (NO) signaling previously associated with iron deficiency [95]. eNOS has a central role in endothelial homeostasis and is generally viewed as serving a beneficial role in lung inflammation [96–98] and ARDS [99, 100]. eNOS uncoupling can lead to generation of reactive oxygen species (ROS) and lung injury [101]. However, this was not indicated in this setting as the ROS signature was reduced in iron deficient mice (Fig. 3b, ROS). A number of other stress responses signatures were also lower in iron deficient mice; specifically, XBP1 (endoplasmic reticulum stress) [102], ATF3 and ATF4 (stress-induced transcription factors) [103–106] (Fig. 3b). In addition, expression of 18 heat shock protein mRNAs was lower (Fig. 3b, DEGs), with Hspa1a and Hspa1b (Hsp70 family members) [66] the most down-regulated DEGs (Supplementary Table 2).

In summary, iron deficient mice at 6 dpi showed modest transcriptional changes when compared to controls. Bioinformatic analyses indicated signatures associated with increased Th1/Th2 ratios, modulated leukocyte expression patterns, and elevated eNOS and reduced stress responses.

### Iron loading vs. control diet; weights, lung viral loads and liver iron levels

Male C57BL/6J mice were fed either a control diet or an iron loading diet for 7 weeks starting at 4 weeks of age (Fig. 4a). The iron loading diet resulted in a small but significant reduction in body weight (mean 1.4 g reduction) at 0 dpi (Fig. 4b). Lung tissue titers at 2 dpi showed a modest, but significant, 0.65 log_10_CCID_50_/g reduction in lung viral titers from mice fed the iron loading diet (Fig. 4c). Nasal turbinate viral titers showed no significant differences at 2 dpi (Supplementary Fig. 2b). Viral lung read counts (from RNA-Seq analysis) also showed a reduction (of 0.39 log_10_), but this did not reach significance (Fig. 4d). Liver iron level analysis confirmed that the iron loading diet had successfully increased iron levels (Fig. 4e).

**Fig. 4.**
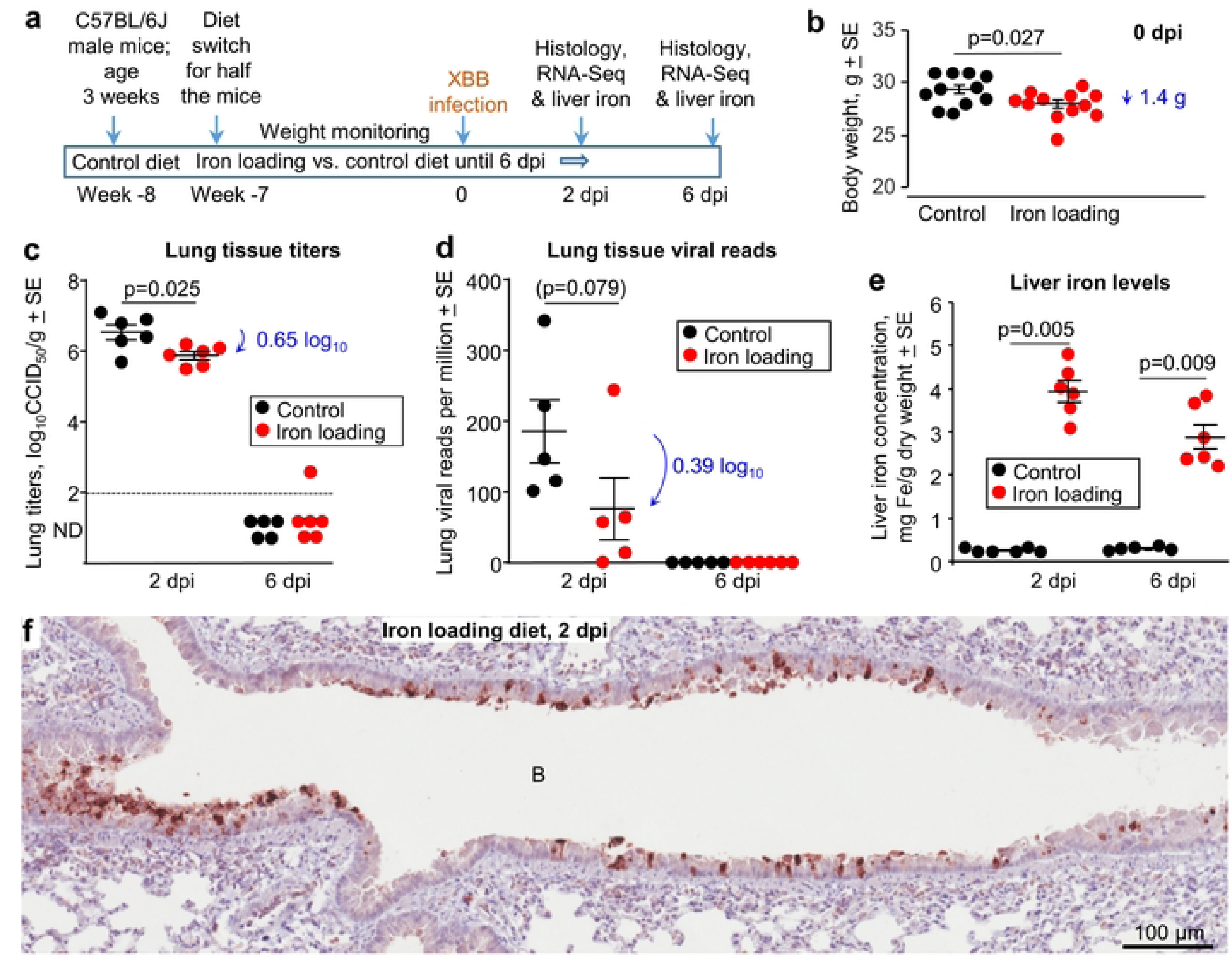
Iron loading vs. control diets; weights, viral loads and iron levels. **a** Time line of experiment. **b** Mouse body weights at 0 dpi, prior to XBB infection. Statistics by t test, n=11/12 mice per group. There was no significant weight changed after infection (Supplementary Fig. 1). **c** Lung tissue XBB virus titers. Limit of detection ≈2 log_10_CCID_50_/g (dashed line); ND – not detected. **d** XBB viral read counts obtained from RNA-Seq of lung tissues. **e** Liver iron levels in the same mice shown in b-d. Statistics by Kolmogorov Smirnov tests. **f** IHC of lung at 2 dpi using the anti-spike monoclonal antibody, for a mouse on the iron loading diet. B – bronchial air space. IHC of lung from a mouse on the control diet is shown in Fig. 1f. Staining of an uninfected lung is shown in Supplementary Fig. 3a.

Thus, the iron loading diet marginally reduced the mean mouse body weight, with lungs showing modest, but significant, viral titer reductions at 2 dpi. The latter is consistent with some [107], but not other [8], *in vitro* studies.

### Iron loading vs. control diet; immunohistochemistry

Whole lung sections were stained with a SARS-CoV-2 specific monoclonal antibody. Staining was primarily associated with the bronchial epithelium, with minimal stained material in the bronchial airway (Fig. 4f). Staining was less abundant (compared with controls, Fig. 1f), consistent with the lower viral load (Fig. 4c). IHC staining of an uninfected mouse lung is shown in Supplementary Fig. 3a.

### Iron loading vs. control diet; histochemistry and lung lesions

Whole lung sections were stained by H&E and scanned slides were examined by a veterinary pathologist. Lung lesions were scored as above, with lesions at 6 dpi emerging to be slightly less severe across the scoring criteria (Supplementary Fig. 4b, Fig. 5a), with the cumulative lesion score significantly lower for iron loaded mice when compared with mice on the control diet (Fig. 5b). This observation likely reflects the lower viral loads in lungs from iron loaded mice (Fig. 4c).

**Fig. 5.**
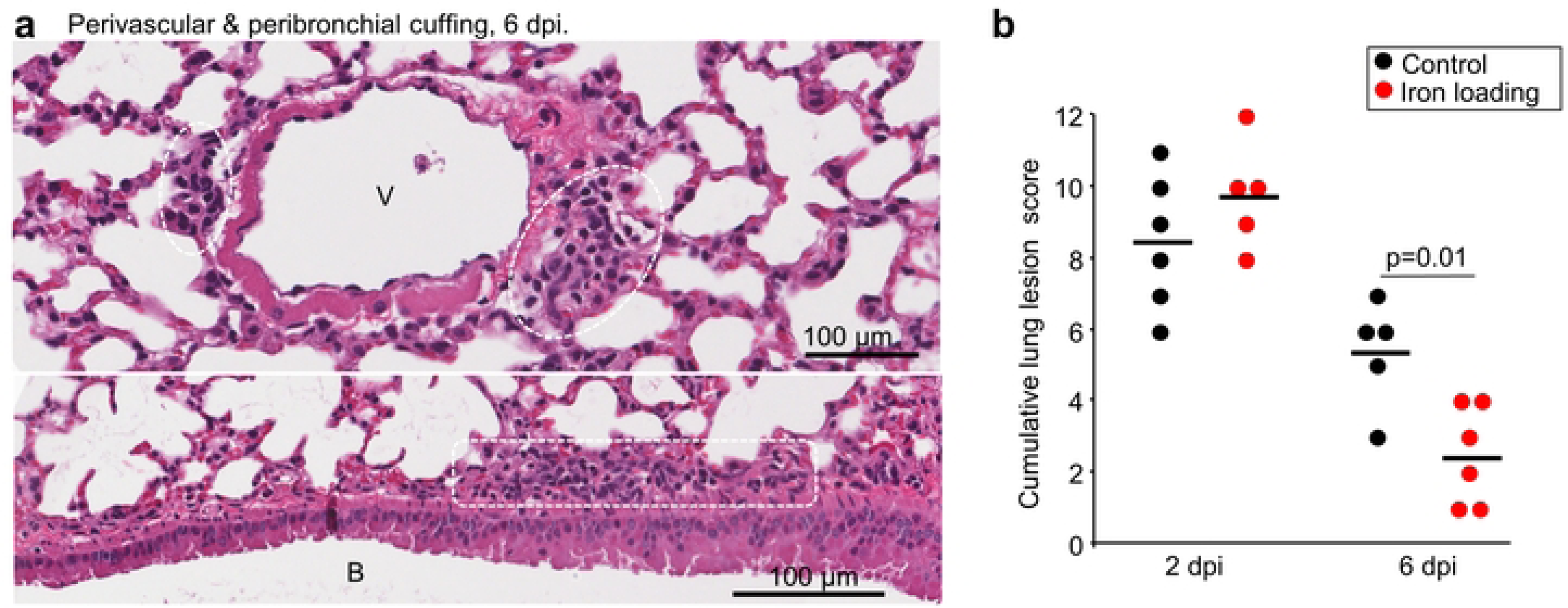
Iron loading vs. control diets; histology. **a** Example of perivascular cuffing (top) and peribronchial cuffing (bottom) at 6 dpi for iron loaded mice. Leukocytes indicated by white dashed box/ovals. V – vascular lumen. B – bronchial lumen. Whole slide score for perivascular/peribronchial cuffing =1. **b** Cumulative lung lesion scores for each mouse, raw data is shown in Supplementary Fig. 4b. Statistics by t test.

White space analysis (Supplementary Fig. 10) and ratios of nuclear (purple) to cytoplasmic (red) staining (Supplementary Fig. 6c) showed no significant differences between infected mice on iron loading vs. control diets.

### Iron loading vs. control diet; lung RNA-Seq at 2 dpi

RNA-Seq analysis of lungs at 2 dpi from mice fed an iron loading vs. control diet identified only 1 DEG, insufficient for meaningful pathway analysis. PC2/PC1 plots are shown in Supplementary Fig. 7b. Full gene lists and bioinformatic analyses are provided in Supplementary Table 3 and are summarized in Fig. 6a (see below).

**Fig. 6.**
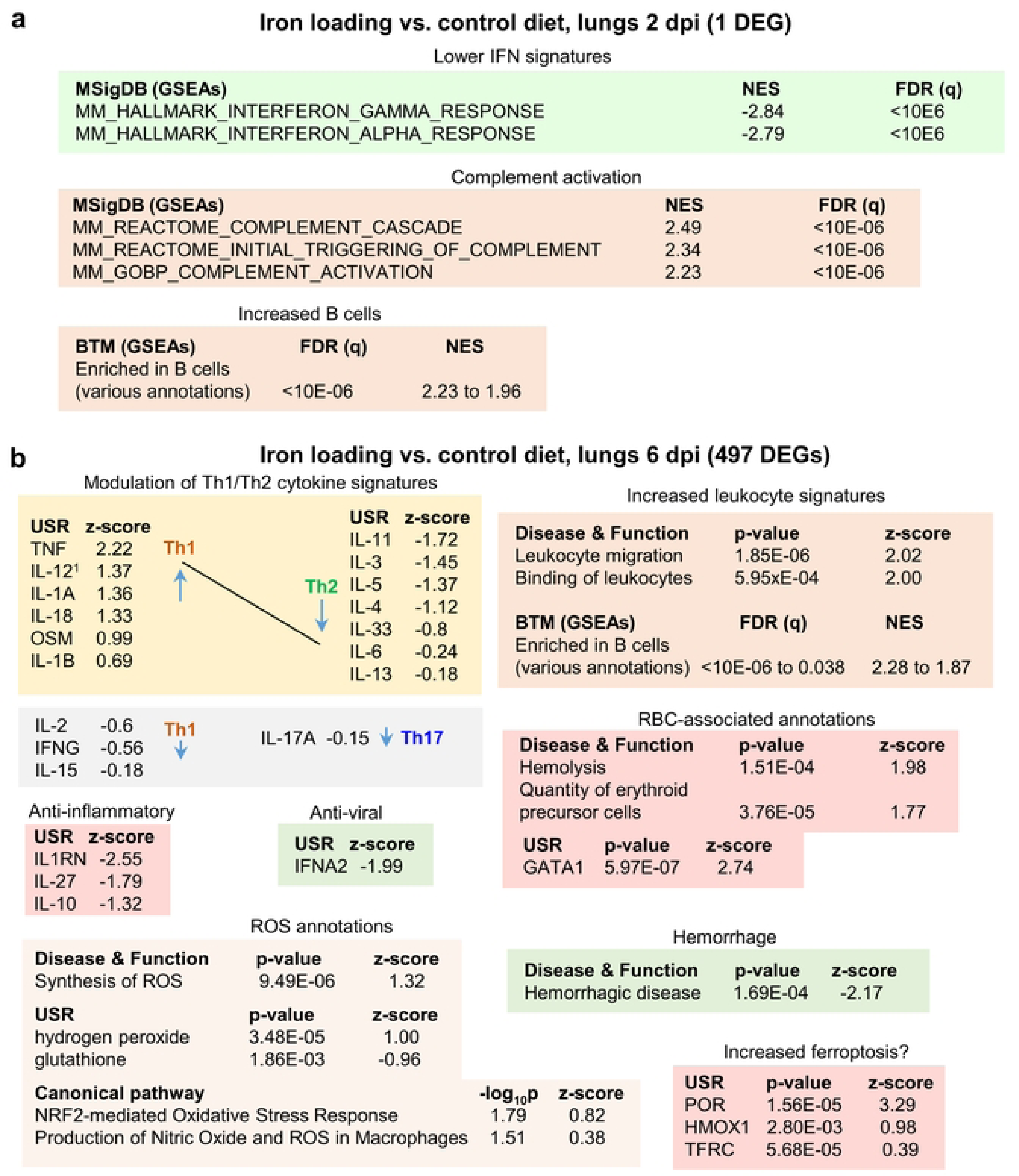
Iron loading vs. control diets; bioinformatic summary. **a** RNA-Seq analysis of lungs from mice fed an iron loading vs. control diet at 2 dpi yielded only 1 DEG, insufficient for pathway analysis. The ‘All genes’ list was interrogated using GSEAs and the gene sets provided by MSigDB. Full data sets are available in Supplementary Table 3. **b** RNA-Seq analysis of lungs from mice fed an iron loading vs. control diet at 6 dpi yielded 497 DEGs. DEGs were analyzed by IPA as in Fig. 3. Full data sets are available in Supplementary Table 4.

GSEAs using the MSigDB gene sets provided a number of IFN annotations with high negative NES values (Fig. 6a). These likely reflect the lower lung viral loads (Fig. 4c,d), with less virus replication in iron loaded mice resulting in less stimulation of IFN responses. MSigDB GSEAs also identified complement activation with high positive NES values (Fig. 6a). SARS-CoV2 is known to activate complement via the lectin [108] and alternative [109] pathways. Complement activation can mediate antiviral effects against SARS-CoV-2 [110], providing a potential explanation for the reduction in viral loads. Iron infusions are reported to trigger complement via the lectin and alternative pathways [111, 112]. One might thus speculate that iron loading reduces the threshold for complement activation during SARS-CoV-2 infection. The one DEGs (Per1) may also be linked to complement activation [113]. However, it should be noted that complement activation primarily involves proteolytic processes that are not directly detectable by RNA-Seq. Complement activation by adaptive immune responses (mediated by IgM/IgG and the classical pathway) comes later during the course of infection, and can also [108] contribute to COVID-19 severity [114]. However, complement activation was not identified in iron loaded mice at 6 dpi (Supplementary Table 4), with virus largely cleared in the current model at this time (Fig. 4c,d).

GSEAs using the BTM gene sets suggested an increased number of B cells infiltrating the infected lungs in iron deficient mice (Fig. 6a).

### Iron loading vs. control diet; lung RNA-Seq at 6 dpi

RNA-Seq analysis of lungs at 6 dpi from mice fed an iron loading vs. a control diet identified 497 DEGs. Fold changes were again modest, with only 2 genes showing a log_2_ fold change >2. Full gene lists and bioinformatics are provided in Supplementary Table 4 and are summarized in Fig. 6b.

Modulation of Th1/Th2 cytokines was again apparent with mostly increased Th1 and reduced Th2 USR cytokine z scores in mice on an iron loading diet (Fig. 6b). This result contrasts with *Salmonella typhimurium* infection of mice on a high iron diet where Th1 responses were inhibited [115]. However, the effects of high iron on the Th1/Th2 balance may be setting specific, with, for instance, iron promoting M1 differentiation of macrophages [21], and lung macrophages playing a central role in COVID-19 inflammation [116]. Several anti-inflammatory USR cytokine signatures provided negative z-scores (Fig. 6b), indicating less anti-inflammatory activity in iron loaded mice. However, the IFNA2 USR signature had a negative z-score (Fig. 6b, Anti-viral), suggesting reduced inflammation, with type I IFNs major drivers of inflammation [117]. A reduced type I IFN signature is consistent with the lower viral loads at 2 dpi (Fig. 4c) and the negative NES for IFN-associated GSEAs at 2 dpi (Fig. 6a).

The annotations associated with increased leukocyte signatures (Fig. 6b) do not reflect significant, histologically observable, increases in leukocytes infiltrates (Fig. 5a, Supplementary Figs. 4b and 6c). This again suggests they are associated with modest infiltrate changes, Th1/Th2 response changes and/or changes in the cell types infiltrating the lungs (see below). A series of BTMs suggested an increase in B cells (Fig. 6b), although no chemokines or chemokine receptors that would readily explain migration of B cells into the lungs were present in the DEG list (Supplementary Table 4).

Iron has often been associated with ROS production [118, 119], and ROS-associated annotations were identified amongst the Canonical pathway annotations. However, z-scores were modest (≤1.3) (Fig. 6b, ROS annotations), perhaps ameliorated by the lower viral loads. Increased ROS in iron overload setting is often ascribe to the Fenton reaction (e.g. [120]), however, the physiological relevance of this reaction *in vivo* is not without controversy [121].

The negative z-score for the hemorrhagic disease annotation (Fig. 6b) likely reflects the reductions in viral load, with lung hemorrhage well documented in COVID-19 mouse models [108, 122]. A number of transcription factor USRs also showed negative z-scores in iron loaded mice (Supplementary Table 4). These include, FOXC1 [123], XBP1 [124], EIF4E [125] and SREBF1 (aka SREBP1) [126], which are induced by SARS-CoV-2 infection. FOXC1 [127], XBP1 [128], and EIF4E [129], as well as TCF3 [130], are also involved in wound repair. Reduced infection (Fig. 4c,d) and/or an ensuing reduced requirement for tissue repair, may explain these negative z-scores.

A number of RBC-associated annotations were identified, including hemolysis (Fig 6b). Although viral infections [80], including SARS-CoV-2 [81], can cause hemolysis, why this should be higher in iron loaded mice is unclear. This may be due to the toxic effects of iron on RBC [131], or is associated with complement activation (Fig. 6a), with complement-mediated hemolysis a well-documented phenomenon [132, 133]. GATA1 is the master regulator of erythropoiesis [134], which might be upregulated to compensate for RBC loss.

POR (NADPH-cytochrome P450 oxidoreductase) was identified as the top scoring USR by z-score (Fig. 6b, Supplementary Table 4). Amongst other functions, POR is involved in the induction of ferroptosis [135]. Ferroptosis is a form of cell death promoted by iron that involves peroxidation of lipids [136], which has been implicated in tissue damage during COVID-19 [137, 138]. Heme oxygenase 1(HMOX1) is a crucial ferroptosis factor [139], and the transferrin receptor (TFRC) is a ferroptosis marker [140], with increased ROS/H_2_0_2_ and reduced glutathione (Fig. 6b) crucial to the process of lipid peroxidation [136]. However, it should be noted that there is no single universal ferroptosis pathway, with many initiators, sensitizers and modulators [141]; hence reliable ferroptosis annotations are often lacking in bioinformatic pathway tools such as IPA.

In summary, iron loaded mice showed modestly lower lung viral loads at 2 dpi, with RNA-Seq indicating modest transcriptional changes. Signatures at 6 dpi in iron loaded mice were associated with a general bias toward increased Th1/Th2 cytokine ratios, hemolysis, ROS and perhaps ferroptosis, but also reduced type I IFN and hemorrhage.

### Dysregulated cell compositions in infected lungs of iron deficient and iron loaded mice

To gain insights into how iron deficiency and iron overload might influence the cellular compositions in the lungs after SARS-CoV-2 infection, cellular deconvolution (SpatialDecon) analysis was undertaken. This used the normalized count matrices (that provide the number of aligned reads for each gene for each mouse) and the gene expression matrices from the Lung mouse cell atlas.

In iron deficient mice at 2 dpi, a significant increase in von Willebrand factor (VwF) positive endothelia cells was identified (Fig. 7a). This observation is perhaps consistent with the increased extravasation annotations (Fig. 3a), as endothelial VwF promotes extravasation [142, 143].

**Fig. 7.**
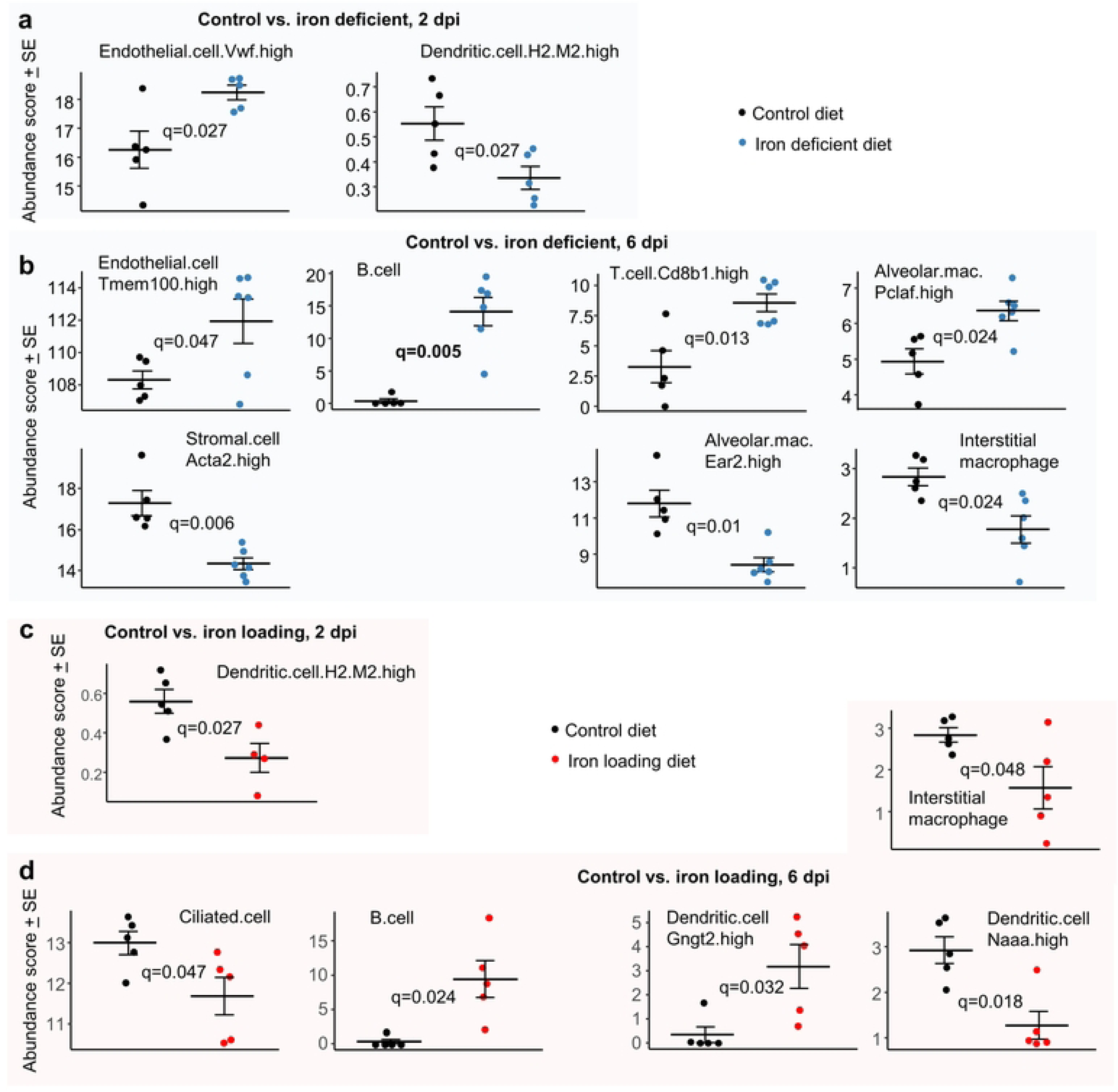
Cellular deconvolution analyses for iron deficient vs. control and iron loading vs. control diets. Relative abundances of cell types were estimated using SpatialDecon and cell-type expression matrices obtained from the NanoString Lung mouse cell atlas. **a** Control vs. iron deficient, 2 dpi. **b** Control vs. iron deficient, 6 dpi. **c**. Control vs. iron loading, 2 dpi. **d**. Control vs. iron loading, 6 dpi. Statistics by t test with FDR correction.

In iron deficient mice at 6 dpi, the increase in endothelial transcriptional signatures (Fig. 7b) is consistent with the increase in the eNOS signaling pathway (Fig. 3b), with eNOS a key survival factor for endothelial cells during inflammation [144]. Stromal cells were less abundant in iron deficient mice; these cells are involved in wound repair [145] and their proliferation may be impaired under iron-deficient conditions [146]. The higher B and T cell abundance scores in iron deficient mice (Fig. 7b) are consistent with the BTM GSEA results (Fig. 3b). Difference in mean abundance scores for B cell represents the largest and most significant (Fig. 7b, q=0.006) observed herein. Lung B and T cells are well described for COVID-19 [147, 148]; however, iron deficiency is generally associated with impaired B and T cell responses [149, 150]. The Interstitial macrophages are generally anti-inflammatory during disease processes [151] and Ear2 is upregulated on alveolar macrophages under Th2 conditions [152], so reduced abundance scores for these two cell types in iron deficient mice (Fig. 7b) may be associated with the elevated Th1/Th2 ratios (Fig. 3b). In contrast, proliferating alveolar macrophages (Pclaf is a proliferation marker) have a higher abundance score in iron deficient mice (Fig. 7b), with these cells known to self-renew and adopt a M1 pro-inflammatory phenotype when exposed to IFNγ or TNF [116].

In iron loaded mice at 2 dpi, only Dendritic.cell.H2.M2.high [153] were identified as significantly less abundant (Fig. 7c), with less infection perhaps resulting in less dendritic cell activation and/or recruitment. In iron loaded mice at 6 dpi, the abundance score for ciliated (epithelial) cells was significantly lower (Fig.7d), with these cells efficiently infected by omicron variants [154]. The lower levels of infection and sloughing in iron loaded mice (Fig. 4c, f) compared with controls (Fig. 1f, top) is consistent with a lower requirement for renewal and thus the reduced (transcription-based) abundance scores. The increased B cell abundance (Fig. 7d) is again consistent with BTM GSEAs (Fig. 6b). DCs expressing N-acylethanolamine acid amidase (Naaa) have a lower abundance score in iron loaded mice (Fig. 7d). These cells are reported to play a proinflammatory role [155], with reduced type I IFN responses (Fig. 6b, IFNA2) perhaps contributing to their lower abundance [156]. Gngt2 is a M1 marker [157] and describes a subgroup of DCs [158], whose specific function has yet to be described.

Cellular deconvolution using expression matrices from the ImmGen cell family (which is based on various tissues, not just lung) further illustrates dysregulation of leukocyte subsets at 6 dpi in both iron deficient and iron loaded mice (Supplementary Fig. 11).

## Discussion

We provide herein detailed comparisons of lung SARS-CoV-2 omicron XBB infection and inflammatory disease in wild-type mice that were fed an iron deficient vs. a control diet, and mice fed an iron loading vs. control diet. In iron deficient and iron loaded mice, viral loads were either not significantly affected or were mildly (≈ 0.6 log_10_), but significantly, reduced in lungs or nasal turbinates (Fig. 4c, Supplementary Fig. 2a). This strongly argues that the effectiveness of the host’s anti-viral SARS-CoV-2 responses was not significantly compromised by altered iron status. Modulation/dysregulation of immune responses by iron deficiency and overload is well described in various setting [159, 160], and was also clearly evident herein (Figs. 3, 6 and 7; Supplementary Fig. 11). However, this led neither to impaired ability to control the virus nor to overtly more severe lung histopathology.

Pleiotropic outcomes of SARS-CoV-2 infections in iron deficient and iron overloaded mice might be envisaged, given the complexity of iron regulation [20], the modulation of iron biomarkers and homeostasis during SARS-CoV-2 infections [14, 23–31], and the robust pro-inflammatory responses associated with SARS-CoV-2 infections [66, 68, 116, 161].

However, perhaps unexpected was the increase in Th1/Th2 ratios and B cell signatures in both iron deficient and iron load mice. This observation is reminiscent of the commonly illustrated U-shaped relationships for iron status. For instance, both iron deficit and iron excess leads to impaired or dysregulated maternal immunity during pregnancy [162]. A similar U shaped relationship is also reported for maternal hemoglobin and preterm birth and ARDS [163]. Serum iron biomarkers and COVID-19 severity also show a similar relationship [23]. B cell responses are also often impaired during iron deficiency [21], with IgM responses also reported to be blunted in iron loaded mice [164]. However, the effect of iron deficiency and overload on adaptive T cell responses would appear to be quite diverse and setting dependent [21, 74, 75, 89, 165, 166]. In addition, in humans early Th1 responses have been associated with protection against severe COVID-19 [167, 168]; however, in mouse models, early pro-inflammatory signatures (2-7 dpi) tend to correlate with pathogenic “cytokine storm” profiles [66, 68]. Infection in humans usually spreads from the upper respiratory track into the lower respiratory track (lungs). This progression is not recapitulated in mice where lung infection requires direct intra-pulmonary inoculation of virus [71, 72, 169]. Whether the increased Th1/Th2 ratios seen herein might be associated with beneficial or detrimental activities is thus debatable.

Perhaps surprising was the wide scale reduction in stress responses in iron deficient mice at 6 dpi (Fig. 3, XBP1, ATF4, ROS). The most dominant of these (by z score) was XBP1, which is associated with the endoplasmic reticulum (ER) unfolded protein response (UPR). This pathway is activated by SARS-CoV-2 infection, promotes viral replication in epithelial cells, and is associated with induction of proinflammatory responses [124, 170]. Why this response is blunted in iron deficient mice may be related to the requirement for iron and heme effectors and binding proteins for Ire1 clustering, a process that lies immediately upstream of XBP1 activation [102]. ATF4 is triggered by PERK, another sensor of ER stress and mediator of the UPR [171]. Similarly, as iron is an important component of ROS-generating enzymes [172], iron deficiency may reduce ROS production capacity in this setting.

A limitation of this study is that we have not investigated the responses over the long-term, with a role for iron status in increasing the severity of long-COVID suggested by several studies [12, 19, 173]. However, whether mouse models [174] can faithfully recapitulate pathological or immunopathological features of human long-COVID remains to be established [175], with underlying co-morbidities [176] clearly absent in genetically identical, specific pathogen free, laboratory mice. We have also not provided insights into the sizable range of co-morbidities that can give rise to anemia or iron overload [53, 177, 178], and how these would affect COVID-19; however, this would constitute a considerable undertaking. For instance, we have not studied the ‘homeostatic iron regulator’ deficient (Hfe^-/-^) mouse for mouse model of hereditary hemochromatosis [179]. A counter rationale for Hfe^-/-^ mouse studies is that compelling evidence for Hfe mutations affecting COVID-19 patient outcomes has yet to emerge [180]. We have also used herein a mouse model of relatively mild disease, as distinct from the more severe K18-hACE2 model. However, the K18-hACE2 model is complicated by early mortality associated with fulminant brain infections [72], which are generally not a feature of human disease [67]. Nevertheless, whether iron status would influence severe lung infection and disease is thus not addressed in our study. Lastly, we have not investigated changes in iron distributions and their immunological consequences over time in different tissues (e.g. lungs, liver, spleen, lymph nodes) [63, 181], primarily as there were no overt detrimental outcomes that would focus such studies.

In conclusion, the current study of iron deficient vs control and iron loaded vs. control SARS-CoV-2 infected mice, finds modest transcriptional changes indicating a range of inflammatory response modulations, but no significant histopathologically detectable disease exacerbations. Some human studies have also failed to find a significant association between iron status and severity of acute COVID-19 [180, 182, 183]. This is not to say that all diseases or conditions that lead to iron deficiency or overload are similarly benign, as they may indicate co-morbidities that can promote COVID-19 severity [184, 185].

## Materials and methods

### Ethics statements and regulatory compliance

Collection of nasal swabs from consented COVID-19 patients was approved by the University of Queensland HREC (2022/HE001492). All mouse work was conducted in accordance with the Australian code for the care and use of animals for scientific purposes (National Health and Medical Research Council, Australia). Mouse work was approved by the QIMR Berghofer MRI Animal Ethics Committee (P3600 and P3535). All infectious SARS-CoV-2 work was conducted in the BioSafety Level 3 (PC3) facility at the QIMR Berghofer MRI (Department of Agriculture, Fisheries and Forestry, certification Q2326 and Office of the Gene Technology Regulator certification 3445). Mice were euthanized using carbon dioxide.

### Iron diet modifications

Male C57BL/6 mice were bred in-house at the QIMR Berghofer MRI animal facility and were held under standard animal house conditions (for details see [70]). Breeding pairs were maintained on standard rodent pellet diet (120 mg/kg iron; Norco Stockfeed, Lismore, Australia). Mice were allowed unlimited access to food and water at all times.

*Iron deficient diet*. Three-week old mice were weaned onto an iron deficient diet based on AIN93G (∼1 mg/kg iron, Specialty Feeds, Glen Forrest, Australia). This iron deficient diet produces a mild to moderate anemia [62, 186]. *Iron loading diet.* Three-week old mice were fed the control diet for one week, after which they were switched to an iron loading diet, consisting of the iron deficient diet supplemented with 0.5% iron as carbonyl iron (Sigma, Product no. C3518). This 1 week delay in switching mice to the iron loading diet is necessary, as moving weanling mice directly onto an iron loading diet dramatically reduces growth rates. *Control diet*. The control diet comprised the aforementioned iron deficient chow supplemented with 50 mg/kg iron as ferric citrate. All mice were maintained on these diets throughout until euthanasia.

### Liver iron quantification

Liver iron levels were assayed by colorimetric assay as described previously [187], with liver slices fixed in formalin to inactivate virus prior to release from PC3.

### The SARS-CoV-2 omicron XBB isolate

The XBB isolate (SARS-CoV-2_UQ01_) was voluntarily donated to the University of Queensland (Brisbane, Australia) by a deidentified adult COVID-19 patient with degree-level education via a self-collected nasopharyngeal swab. The patient provided written consent [65, 67]. The isolate was initially grown on Vero E6-TMPRSS2 cells [188]. The isolate is XBB.1.9.2.1.4 (Pango EG.1.4), a recombinant of BA.2.10.1 and BA.2.75; sequence deposited as hCoV-19/Australia/UQ01/2023; GISAID EPI_ISL_17784860. XBB viral stocks were propagated in Vero E6 cells [71], and were titered using CCID_50_ assays [189]. Medium was checked for endotoxin [190] and cultures for mycoplasma (MycoAlert, Lonza).

### Mice infection and monitoring

Mice received intrapulmonary infections delivered via the intranasal route with 5×10^4^ CCID_50_ of virus in 50 μl RPMI 1640 whilst under light anesthesia as described [72]. Mice were weighed and monitored as described [67, 72].

Mice were euthanized using CO_2_, lungs were removed, with the left lung fixed in formalin for histology, the right lung inferior lobe placed in RNAlater for RNA-Seq and the remaining lobes used for tissue titers determination by CCID_50_ assays using Vero E6 cells as described [71, 72].

### CCID_50_ assays

Tissue titers were determined as described [71]. Briefly, 5-fold serial dilutions of clarified tissue homogenates were applied in duplicates to Vero E6 cells in 96 well plates. After 6 days cytopathic effects were observed by inverted light microscope. The virus titer was determined by the method of Spearman and Karber; an Excel sheet is available at https://www.klinikum.uni-heidelberg.de/zentrum-fuer-infektiologie/molecular-virology/welcome/downloads.

### Immunohistochemistry

Immunohistochemistry was undertaken using the anti-SARS-CoV-2 spike monoclonal antibody, SCV2-1E8 as described [69], except that the monoclonal (IgG2a) was purified using Protein A affinity chromatography and applied to sections at 4 µg/ml for 1 hr.

### Histology

Lungs were fixed in 10% formalin, embedded in paraffin, and sections stained with H&E (Sigma Aldrich). Slides were scanned using Aperio AT Turbo (Aperio, Vista, CA, USA). Quantitation of white space in scanned images of H&E stained lung parenchyma (with areas greater than ≈100 μm set as a threshold) was undertaken using PixelClassifierTools in QuPath v0.3.2 [191], and provides an approximate measure of lung consolidation [72].

Scanned H&E stained whole lung sections were analyzed by Aperio Positive Pixel Count Algorithm (Leica Biosystems) to generate nuclear (strong purple staining) over cytoplasmic (total red staining) pixel count ratios, providing an approximate measure of leukocyte infiltration [72, 73].

All H&E stained whole lung sections were scanned and .svs files examined by a trained European board-certified veterinary pathologist using Qu-Path (v 0.5.1). Lung lesions were scored using 6 criteria. Emphysema was scored; 0=no lesion, 1=dilated and coalescent alveoli, 2=“bullae” in the parenchyma. Bronchial epithelium damage was score; 0= no lesion, 1=small clusters of necrotic epithelial cells, 2=scattered foci of epithelial degeneration with layer architecture partial loss, 3=focal complete epithelial loss. Bronchial content was scored; 0= empty lumen; 1=presence of a small amount of material; 2=partial obliteration; 3=complete occlusion. Vascular wall changes were scored; 0= no lesion, 1=leukostasis, 2=focal wall damages (including leukocytoclasis), 3=transmural vessel wall alteration and/or vascular lumen obliteration. Perivascular edema was scored; 0=no lesion, 1=focal mild edema, 2=extended marked edema with lymphoid vessel dilatation.

Peribronchial/perivascular cuffing was scored; 0= no lesion; 1=focal inflammatory cell infiltration; 2=circumferential inflammatory cell infiltration, 3=coalescing inflammatory cell infiltration between bronchi and vessels. A total cumulative score was then calculated by summing all 6 parameter scores for each mouse (range 0 to 16).

### RNA-Seq and bioinformatics

RNA-Seq (Illumina Nextseq 2000 platform generating 75 bp paired end reads) and bioinformatics was undertaken as described [66, 68]. Mean quality scores were above Q20 for all samples. Mouse RNA-Seq reads were aligned to a combined mouse (GRCm39, version M27) and SARS-CoV-2 BA.5 reference genome [67] using STAR aligner. Viral read counts were generated using Samtools v1.16. RSEM v1.3.1 was used to generate expected counts for host genes. EdgeR was then used to generate TMM normalized count matrices, with a separate count matrix generated for iron deficient vs. control and iron loaded vs. control. Differentially expressed genes were identified using EdgeR using a FDR cut-off of 0.05.

Pathway analyses were performed using host DEGs and Ingenuity Pathway Analysis (IPA, v84978992) (QIAGEN), which provides Canonical pathways, Up-Stream Regulators (USR) and Diseases or Functions features as described [66, 72]. Annotations without z scores or with significance (q or p) below 0.05 were removed.

Gene Set Enrichment Analyses (GSEAs) were undertaken using GSEA v4.1.0 with gene sets provided in MSigDB (≈ 45,000 gene sets) and in the Blood Transcription Modules [192], and gene lists ranked by log_2_ fold-change. Relative abundances of cell types were estimated in R v4.1.0 from TMM normalized RSEM count matrices using SpatialDecon v1.4.3 [193] and cell-type expression matrices obtained from the Lung mouse cell atlas (https://github.com/Nanostring-Biostats/CellProfileLibrary/blob/master/Mouse/Adult/Lung_MCA.RData) and the ImmGen cell family (https://github.com/Nanostring-Biostats/CellProfileLibrary/blob/master/Mouse/Adult/ImmuneAtlas_ImmGen_cellFamily.R <u>Data</u>). Statistics were undertaken by t tests with False Discovery Rate corrections using the Benjamini-Hochberg method (q).

### Statistics

The t-test was used if the difference in variances was <4 fold, skewness was > - 2 and kurtosis was <2. The t test significance and variance were determined using Microsoft Excel.

Skewness and kurtosis were determined using IBM SPSS Statistics for Windows v19.0 (IBM Corp., Armonk, NY, USA). Otherwise, the non-parametric Kolmogorov-Smirnov exact test was performed using GraphPad Prism 10.

## Supporting information

The manuscript is accompanied by 4 Supplementary Tables and 11 Supplementary Figures.

## Acknowledgments

The authors thank the following QIMRB staff; Dr. I. Anraku for management of the PC3 facility at QIMR Berghofer MRI, Dr. Viviana Lutzky for proof reading, Dr Crystal Chang for histology services, the animal house staff for mouse breeding and agistment, and Dr. Gunter Hartel for assistance with statistics.

## Author Contributions

### Conceptualization

AS, DF, JH

### Data curation

AS, CRB

### Formal analysis

TL, CRB

### Funding Acquisition

AS, DJR

### Investigation

AC, KY, BT, WN, SLH.

### Methodology

CRB, KY, DJR, CRB, SLH

### Project Administration

AS, DJR

### Resources

AS, DJR

### Software

CRB

### Supervision

AS, CRB, DJR, WN

### Validation

AS, CRB

### Visualization

AS, TL

### Writing – original draft

AS, AC

### Writing – review & editing

AS, AC, DF, JH, TL

## Data availability statement

All data is provided in the manuscript and accompanying supplementary files. Raw sequencing data (fastq files) generated for this publication for RNA-Seq have been deposited in the NCBI SRA, BioProject: PRJNA1102925 and are publicly available.

